# Prediction and Expression Analysis of Deleterious Nonsynonymous SNPs of Arabidopsis ACD11 Gene by Combining Computational Algorithms and Molecular Docking Approach

**DOI:** 10.1101/2021.10.08.463616

**Authors:** Mahmudul Hasan Rifat, Jamil Ahmed, Milad Ahmed, Foeaz Ahmed, Airin Gulsan, Mahmudul Hasan

## Abstract

Accelerated cell death 11 (ACD11) is an autoimmune gene that suppresses pathogen infection in plants by preventing plant cells from becoming infected by any pathogen. This gene is widely known for growth inhibition, premature leaf chlorosis, and defense-related programmed cell death (PCD) in seedlings before flowering in *Arabidopsis* plant. Specific amino acid changes in the ACD11 protein’s highly conserved domains are linked to autoimmune symptoms including constitutive defensive responses and necrosis without pathogen awareness. The molecular aspect of the aberrant activity of the ACD11 protein is difficult to ascertain. The purpose of our study was to find the most deleterious mutation position in the ACD11 protein and correlate them with their abnormal expression pattern. Using several computational methods, we discovered PCD vulnerable single nucleotide polymorphisms (SNPs) in ACD11. We analysed the RNA-Seq data, identified the detrimental nonsynonymous SNPs (nsSNP), built genetically mutated protein structures and used molecular docking to assess the impact of mutation. Our results demonstrated that the A15T and A39D variations in the GLTP domain were likely to be extremely detrimental mutations that inhibit the expression of the ACD11 protein domain by destabilizing its composition, as well as disrupt its catalytic effectiveness. When compared to the A15T mutant, the A39D mutant was more likely to destabilize the protein structure. In conclusion, these mutants can aid in the better understanding of the vast pool of PCD susceptibilities connected to ACD11 gene GLTP domain activation.

## 1. Introduction

Plant possesses an immune system to defend themselves during interactions with pathogen and many component play significant roles in this defense mechanism. For the sake of defense response, programmed cell death (PCD) or apoptosis occurs, and it occurs during various developmental processes like mature pollen stage, visible stage of two to twelve leaves, stage of germinated pollen, flowering stage, stage of mature plant embryo, as well as stage of petal differentiation and expansion, bilateral cotyledonary, globular stage of plant embryo and finally in vascular leaf senescent stage of plants [1–4]. In *Arabidopsis*, during infection the accelerated-cell-death11 (ACD11) response to salicylic acid (SA) resulting PCD and cease pathogen infection. Moreover, ACD11 also performs a role in ceramide transport as a ceramide-1-phosphate transfer protein (second messengers in apoptosis) and as a regulator of phytoceramide. In addition, it also acts in intermembrane lipid transfer and represent itself as sphingosine transmembrane transporter which also response to apoptosis [5–7]. Thus, in *Arabidopsis* ACD11 gene is associated with multiple function starting from plant development to immune response against any stress or pathogen.

Mutant of the ACD11 provides a genetic model for studying immune response activation in *Arabidopsis*. As it is proved that ACD11 is associated with sphingolipid, so any disruption in this gene may cause PCD. For example, previous study revealed that this lethal, recessive, mutant gene could activate immune response and PCD in the absence of pathogen attack or any stress condition that knockout ACD11 mutant, reveals PCD which is SA-dependent [8–9]. In *Drosophila*, disruption of sphingolipid metabolism cause apoptosis which is associated to reproductive defects [10]. Another study hypothesized that the non-existence of ACD11 may be perceives by the agnate nucleotide-binding as well as leucine-rich repeat (NB-LRR) protein, which subsequently triggers PCD [11].

Single nucleotide polymorphisms (SNPs) are the most common type of variation which is abundantly found. In the human genome, SNPs occurs at a frequency of approximately every 100 to 300 base pairs. In short, SNP represents replace or change of a single nucleotide which is called DNA building block. For instance, in a stretch of DNA, SNP may replace the nucleotide cytosine (C) with the nucleotide thymine (T) that is a single nucleotide [12]. Maximum SNPs are synonymous and thus neutral allelic variants. However, main targets of SNP research mainly focus on either the identification of functional SNPs or non-synonymous SNP which is responsible for crop improvement, bringing complex traits and diseases in plants as well as in animals. In crop improvement, single nucleotide polymorphisms (SNPs) is considered as a great source of genetic variations which is not lethal and is associated with cold resistance, draught resistance and disease resistance such as blight, bacterial canker etc. [13–15]. A study in Tea showed that, current *Camellia sinensis* and its wild relatives has genetic divergence which is revealed using genome-wide SNPs from RAD sequencing [16]. In rice, genetic diversity was analyzed using SNP based approaches and revealed important alleles associated with seed size in rice [17].

However, sometimes deleterious nonsynonymous SNPs could have lethal effect on plant and could be dangerous for crops especially when it occurs within a regulatory region of gene. These non-synonymous SNP have the ability to alter the DNA sequence which will lead to disruption in the amino acid sequence of a protein resulting in a biological change in any individual. This is because SNP induces functional impact in protein, for example in protein stability. Therefore, the interaction with other proteins is hampered [18–19]. This deleterious effect could be predicted in *A. thaliana* and likely in other plant species using bioinformatics tools. A previous study identified the SNP diversity in recently cultivated tomato and wild type tomato species by using computational tools [20–21]. In addition, another study revealed that in other eukaryotes, CYP1A1 gene, belonging to the cytochrome P450 family, induces production of polycyclic aromatic hydrocarbon in the lungs and resulting in cardiovascular pathologies, cancer, and diabetes like diseases. SNP rate was higher in this gene and those diseases were predicted using a systematic *in silico* approach. Moreover, CYP11B2 gene undergoes SNP which was also been predicted using computational approaches [22–23]. Thus, there are many bioinformatics tools are being used for predicting SNP in both plant and animal.

Bioinformatics tools make the research easier, resourceful and well ordered. Nowadays, whole genome sequencing of many plants, animals, and microorganisms has revealed polymorphism, gene sequence variation, genetic marker, SNP and so on. But this big data analysis required computational approaches for predicting these in short time and for saving resources before going for wet lab practices. Moreover, *in silico* SNP analysis also facilitate the research and predict the most deleterious and damaging SNPs [24–25]. For example, mutated structure of protein or motif binding may be changed because of SNP, but it has direct correlation with gene expression and variation which could be predicted using computational approaches. Either the SNP is synonymous or nonsynonymous, lethal or not, and have any serious impact on plant or not, all these could be predicted using computational approach [26–29].

Here, we focus on predicting the deleterious nonsynonymous SNPs of Arabidopsis ACD11 gene using computational approaches. Previous research suggests that this kind of study is possible, and SNP diversity with its effects are already identified in recent cultivated tomato and wild tomato species following molecular simulations [30]. As of now, ACD 11 is not well studied and SNP in this gene could be lethal for *Arabidopsis* which may induced PCD in the absence of infection resulting loss of plant and these reasons make us curious, inquisitive to work with this gene.

## 2. Methods

### 2.1 Acquisition of sequences and retrieval of protein crystal structure

All the data of the ACD11 gene were retrieved from various web-based data resources such as The Arabidopsis Information Resource (TAIR) (www.arabidopsis.org), Ensemble Plant (https://plants.ensembl.org/index.html), and Nucleotide and Protein database of National Center for Biotechnology Information (https://www.ncbi.nlm.nih.gov/) and the amino acid sequence (FASTA format) of the reference protein was obtained from the UniProt database (ID-O64587) (https://www.uniprot.org/). Protein sequences and the Protein Deformylases (PDF) corresponding structures were retrieved from the RCSB (Research Collaboratory for Structural Bioinformatics), Protein Data Bank (PDB) (http://www.rcsb.org/pdb/), and a global repository for structural data on biological macromolecules [31]. The protein model with PDB ID: 4NT2 was chosen for the subsequent research work. The PDBSum (http://www.ebi.ac.uk/thorntonsrv/databases/cgbin/pdbsum/GetPage.pl?pdbcode=index.html) was used to gather several key structural information deposited at the PDB.

### 2.2 Analyzing cellular localization and gene expression ACD11 gene in plant physiology

ePlant (http://bar.utoronto.ca/eplant) offers an analytic visualization of multiple levels of *Arabidopsis thaliana* data by connecting a number of freely accessible web services. The tool downloads genome, proteome, interactome, transcriptome, and 3D molecular structure data for the gene(s) or the gene products of interest in a form of conceptual hierarchy [32]. The ePlant tool was used for the single-cell analysis and biotic stress expression including the environmental, pathological and entomological aspects of the ACD11 gene. The SUB cellular location database for Arabidopsis proteins (SUBA4, http://suba.live) is a detailed collection of published data sets that have been manually curated. It uses a list of *Arabidopsis* gene identifiers to provide relative compartmental protein abundances and proximity relationship analysis of protein-protein interaction (PPI) and co-expression partners [33]. The SUBA4 database was employed to generate a confidence score for each distinct subcellular compartment or region, with experimentally-determined localizations being weighted five times more than the predicted ones. The expression of the ACD11 gene in different stages of the plant life cycle was investigated using RNA-Seq and Affymetrix microarray ATH1 GeneChips (Affymetrix, Santa Clara, CA, USA) data. The ePlant (http://bar.utoronto.ca/eplant) and the eFP-Seq Browser (https://bar.utoronto.ca/eFP-Seq_Browser/) allows exploring RNA-seq-based gene expression levels for the gene of interest [34]. GEO Affymetrix microarray data (https://www.ncbi.nlm.nih.gov/geo/) and NASCArrays Information (http://arabidopsis.info/affy) tools was utilized in the process. The RNA-seq profiling data of the *Arabidopsis thaliana* were generated by developmental transcriptome. Total RNA was extracted with RNeasy Plant Kit and Illumina cDNA libraries were generated using the respective manufacturer’s protocols. cDNA was then sequenced using Illumina HiSeq2000 with a 50bp read length [35]. The read data are publicly available in NCBI’s Sequence Read Archive under the BioProject (GEO accession: PRJNA314076). Reads were then aligned to the reference TAIR10 genome using TopHat [36–37]. Reads per gene were counted with Python script using functions from the HTSeq package [38]. The developmental data were taken from ePlant server [39–40]. Gene expression data generated by the Affymetrix ATH1 array [41] and were normalized by the GCOS (GeneChip Operating Software) method [42] and the analysis parameter of TGT value was 100. Most tissues were sampled in triplicate. The *Arabidopsis* ATH1 Genome Array, designed in collaboration with The Institute for Genomic Research (TIGR), contains more than 22,500 probe sets representing approximately 24,000 gene sequences on a single array. The R Project for Statistical Computing (https://www.R-project.org/) provides a wide variety of statistical and graphical techniques, and is highly extensible. Based on the microarray data, the R programming is used to scrutinize the degree to which ACD11 gene expression varies during several stages of the plant growth.

### 2.3 Tissue specific expression of ACD11 gene

Using the ePlant tools, tissue specific expression of the ACD11 gene was examined, including gene expression in the embryo developmental stage, the stem epidermis and vascular bundle area, micro gametogenesis, stigma, and ovaries. The gene expression analysis data was obtained from the ePlant server and NASCAffimatrix microarray data [43] (http://bar.utoronto.ca/NASCArrays/index.php), and all of the tissue-specific RNA-Seq data came from separate experiments. Wild-type Col-0 ecotype *Arabidopsis thaliana* plants were used to obtain embryo developmental expression, epidermis expression, and xylem and cork expression data. Laser capture micro dissection was used to generate embryo developmental data from plant embryos maintained under 16/8-hour light/dark conditions. Manual dissection with forceps was used to extract epidermal expression data from 3 cm sections of the top and bottom of the 10-11 cm long primary stems of treated plots under 18/6-hour light/dark conditions at 100 mEinstein, 22°C, and 50%-70% relative humidity [44]. Secondary thickened hypocotyl was created by continuous removal of the inflorescence stem for 10 weeks, and the plants were maintained under continuous light conditions at 22°C to obtain the xylem and cork expression data (https://www.ebi.ac.uk/arrayexpress/experiments/E-GEDO-6151/samples/?s_page=1%20&s_page%20size=25). Landsberg erecta (Ler) ecotype *Arabidopsis thaliana* plant flowers were utilized to acquire micro gametogenesis, stigma, and ovary expression data, same as they were for embryo development and vascular bundle area. After emasculating stage 8 buds of flowers, data on stigma and ovary tissue expression was produced from isolated pistils. Pistils were collected and frozen in liquid N2 after one day of growth, stigmas were detached from pistils with superfine scissors, and the remaining ovaries were put in separate tubes on dry ice until collection was complete [45]. Pollen from *Arabidopsis* plants in the 5^th^ to 10^th^ development stages, cultivated under 16/8-hour light/dark conditions at 21°C, was used to produce micro gametogenesis expression data [46]. All the tissue specific RNA was isolated and hybridized to the ATH1 GeneChip. Microarray Suite version 5.0 (MAS 5.0) was used to analyze the data, with Affymetrix default analysis settings and global scaling (TGT 100) as the normalization method.

### 2.4 Expression analysis of ACD11 gene in various stress condition

Using the eplant server expression analysis tool, the ACD11 gene expression was examined under abiotic conditions such as heat, cold, osmotic, salt, drought, wounding, and other environmental variables. Using the same browsing tool, the pathological and entomological aspect of the ACD11 gene was also scrutinized. All the abiotic and biotic expression data was generated form wild-type Columbia-0 ecotype *Arabidopsis thaliana* plants and all of the pathological expression data was collected in triplicates from half and full infiltrated leaves. The pathological gene expression data was generated form 5-week-old plants where half and full portion of a plant leaf getting infected with Phytophthora infestans respectively. Plants were grown at 22°C with a light/dark cycle of 8/16 hours and bacterial infiltration performed with 10-8 cfu/ml in 10 mM MgCl_2_ (GEO accession: GSE5616). The entomological data was gathered from an *Arabidopsis* plant that was cultivated in soil at 20°C with a 16/8 hours of light/dark cycle for 3-4 weeks before being cultured with *Myzus persicaere* (apterous aphids) in clip cages and collected the leaves after 8 hours (GEO accession: GSM157299). Then RNA was isolated and hybridized to the ATH1 GeneChip [47]. Aside from the biotic stress, the abiotic stress expression study was performed at 18-day-old plants that were cultivated under long-day conditions of 16/8 hours of light/dark, 24°C, 50% humidity, and 150 Einstein/cm^2^ sec light intensity and this expression analysis was a part of the AtGenExpress project (https://www.arabidopsis.org/portals/expression/microarray/ATGenExpress.jsp). The data for cold and heat stress were collected in a 4°C crushed ice-cold chamber and 3 hours at 38°C followed by recovery at 25°C, respectively. Punctuation of the leaves with three successive applications of a custom-made pin-tool with 16 needles was used to collect wounding expression data. Similar to other experiments, the osmotic, salt, drought and oxidative stress also performed by 300 mM Mannitol, 150mM NaCl and rafts were exposed to the air stream for 15 min and 10 uM Methyl viologen accordingly [48]. All the tissue specific RNA was isolated and hybridized to the ATH1 GeneChip. Microarray Suite version 5.0 (MAS 5.0) was used to analyze the data, with Affymetrix default analysis settings and global scaling (TGT 100) as the normalization method.

### 2.5 Single nucleotide polymorphism (SNP) annotation in ACD11 genes

The 1001 Genomes Project (https://tools.1001genomes.org/polymorph/) has already released a complete investigation of 1135 *Arabidopsis thaliana* genomes, with the goal of annotating them with transcriptome and epigenome data, is a powerful resource for polymorphism study in the reference plant. The nsSNP data of the ACD11 gene were extracted from the 1001 Genome project and considered for further analysis. Beside this, the Ensemble Plant web server presents the variant table (https://plants.ensembl.org/Arabidopsis_thaliana/Tools/VEP) which analyze the 1001 genome project data and predict their effects.

### 2.6 Determination of functional SNPs in coding regions

Sorting Intolerant From Tolerant (SIFT) was used to see how each amino acid substitution affects protein function in order to distinguish between tolerant and intolerant coding mutations. It aligns data at each position in the query sequence to predict damaging SNPs based on the degree of conserved amino acid residues to the closely related sequences. Substitutions with probabilities less than or equal to 0.05 are considered intolerant or deleterious, while those with probabilities greater than or equal to 0.05 are expected to be tolerated [49–50]. Protein Analysis through Evolutionary Relationships (PANTHER) predicts pathogenic coding variants based on evolutionary conservation of amino acids. It uses an alignment of evolutionarily linked proteins to determine how long the current state of a given amino acid has been preserved in its ancestors. The higher the risk of functional consequences, the longer the retention period [51]. The Protein Variation Effect Analyzer (PROVEAN) is a sequence based prediction tool that was employed to predict the damaging effect of nsSNPs in the ACD11 gene. The tool utilizes delta alignment scores that measures the change in sequence similarity of a protein before and after the introduction of an amino acid variation. An equal score or below the threshold of −2.5 indicates deleterious nsSNP alignment [52]. PolyPhen2, examines the protein sequence and replacement of amino acids in protein sequence to predict the structural and functional influence on the protein. If any amino acid alteration or a mutation is detected in protein sequence, it classifies SNPs as possibly damaging (probabilistic score >0.15), probably damaging (probabilistic score >0.85), and benign (remaining) [53]. Furthermore, PolyPhen2 calculates the position-specific independent count (PSIC) score for each variant in protein. The difference of PSIC score between variants indicates that the functional influence of mutants on protein function directly [54]. Using the PolyPhen2, Panther Server, and PROVEAN algorithms, the effects of SIFT were investigated further by looking at the influence of nsSNPs on the structure and function of the protein.

### 2.7 Identification of potential domains in ACD11

A number of servers and tools were utilized for understanding the available protein domains of ACD11 protein and its associated protein superfamily and subfamily. To get an insight into the domain locations of the ACD11 gene and the positions of the possible superfamily domains, the servers Gene3D (1.10.3520.10) and Superfamily Server (SSF110004) were used. Gene3D (http://gene3d.biochem.ucl.ac.uk) is a database that contains protein domain assignments for sequences from all of the major sequencing databases. Domains are predicted using a library of representative profile HMMs generated from CATH super families or directly mapped from structures in the CATH database. The server facilitates complicated molecular function, structure, and evolution connections [55]. SUPERFAMILY is a structural and functional annotation database for all proteins and genomes. This service annotates structural protein domains at the SCOP superfamily level using a set of hidden Markov models. A superfamily is a collection of domains with a shared evolutionary history [56]. Furthermore, PANTHER (PTHR10219) and Pfam (PF08718) were used to investigate the protein subfamily of the ACD11 protein. The PANTHER (Protein Analysis through Evolutionary Relationships) Classification System was created to help high-throughput analysis by classifying proteins (and their genes). Proteins are divided into families and subfamilies. Pfam is a protein family and domain database that is frequently used to evaluate new genomes and metagenomes, as well as to drive experimental work on specific proteins and systems. A seed alignment for each Pfam family comprises a representative collection of sequences for the entry [57].

### 2.8 Homology modelling, validation and molecular docking study

Three-dimensional protein structure models can be built by homology modeling which utilizes experimentally determined structures of related family members as templates. On the basis of a sequence alignment between the target protein and the template structure, a three-dimensional model for the target protein is generated [58]. I-TASSER is an online platform which implements the TASSER-based algorithms and helps to predict the structure of a given protein. In this study, we used I-TASSER for A15T and A39D mutation modeling and then carried out the mutational protein modeling [59]. Then the effects of A15T and A39D mutations in the native protein structure were visualized by Pymol. Next, we considered the ERRAT [60], varify3D [61], [62] and PROCHECK [63] programs to determine and validate the structural stability and residue quality of mutant and native protein. To assess the impact of a particular mutation on the local and global environment of ACD11 protein structure, we have calculated van der Waals, hydrogen bonding, electrostatic and hydrophobic interactions in ACD11 mutant using Arpeggio web server [64]. Furthermore, molecular docking was performed by AutoDock Vina software which allows the binding of the mutant ACD11 structure with the entire surface of the native ACD11 protein. Finally, the docked complexes were analyzed and visualized by Pymol [65].

## 3. Results

### 3.1 Acquisition of sequences and retrieval of protein crystal structure

We utilized the ACD11 gene’s genomic sequence, which is found on chromosome 2 between 14,629,986 and 14,632,082 kb of forward strand, has 4 exons and 3 introns. This gene codes for a glycolipid transfer protein (GLTP) family protein with a 1363bp (NM 129023.5) mRNA that translates into a 206 amino-acid protein (NP 181016.1) (**S1 Table**). This protein contains just one chain in its crystal structure (PDB 4NT2), with 14 helices, 30 helix-helix interactions, and 4 beta turns. This protein contains 5 SO_4_ (Sulphur-di-oxide) ion contacts, 2 SPU (Sphingosylphosphorylcholine), and 2 EDO (Ethylene glycol) ligand interactions and also interacts with the proteins BPA1, PRA1F2, and PRA1F3. The molecular weight of this protein is 22681.60 Da, The IEP (isoelectric point) value is 8.47 and the GRAVY (grand average of hydropathy) value of 0.05 (**S2 Table**).

### 3.2 Analyzing cellular localization and gene expression of ACD11 gene in plant physiology

#### 3.2.1 Cellular localization

*In vitro*, the ACD11 protein transfers sphingosine, a glycolipid precursor, through membranes [66]. As a result, we examined gene expression at the cellular level. The output clearly explained that the ACD11 is a transmembrane protein as this gene is strongly expressed in the cell membrane region. Aside from this location, the ACD11 gene had been found to be expressed in a variety of ways across the cell, apart from the vacuole. In the cytosol and mitochondrion, the ACD11 gene is abundantly expressed. It also had a medium degree of expression in the nucleus and plastid, and a very low level of expression in the endoplasmic reticulum, golgi, peroxisome, and extracellular location. (**S1 Fig and S3 Table**)

#### 3.2.2 RNA-Seq data and developmental transcriptome expression

As the ACD11 gene causes rapid cell death of plants in different abiotic and biotic stress conditions, we further analyzed this gene expression in the different stages of the plant life cycle using RNA-Seq and Affymetrix microarray data to find out when and where this gene is expressed highly in normal condition. From the RNA-Seq analysis data, it is clear that the ACD11 gene is strongly expressed in the mature leaf, first stage of germinating seeds, leaf petiole of the mature leaf, and petals of the mature flower. In the hypocotyl of seedling, leaf lamina of mature leaf, carpel of the mature flower, senescent internodes, and in the root apex, the ACD11 gene is expressed moderately. Apart from these locations, the ACD11 gene is poorly expressed in seeds from the senescent silique, pod of the silique with seed and without seed condition, dry seed and leaf petiole of the young leaf (**Fig 1A**) (**S4 Table**). According to developmental transcriptome data, the ACD11 gene dramatically increases its expression in mature pollen, cauline leaf, second internode, 24 hour imbed seeds, and other floral components. This gene is expressed moderately in the cotyledon, distal half of the leaf, sepals, petals, rosette leaf, and root part. In seeds with and without silique, vegetative rosette, and 9^th^ to 12^th^ flower stage, the lowest expression is anticipated (**Fig 1B)** (**S5 Table**).

**Fig 1.**
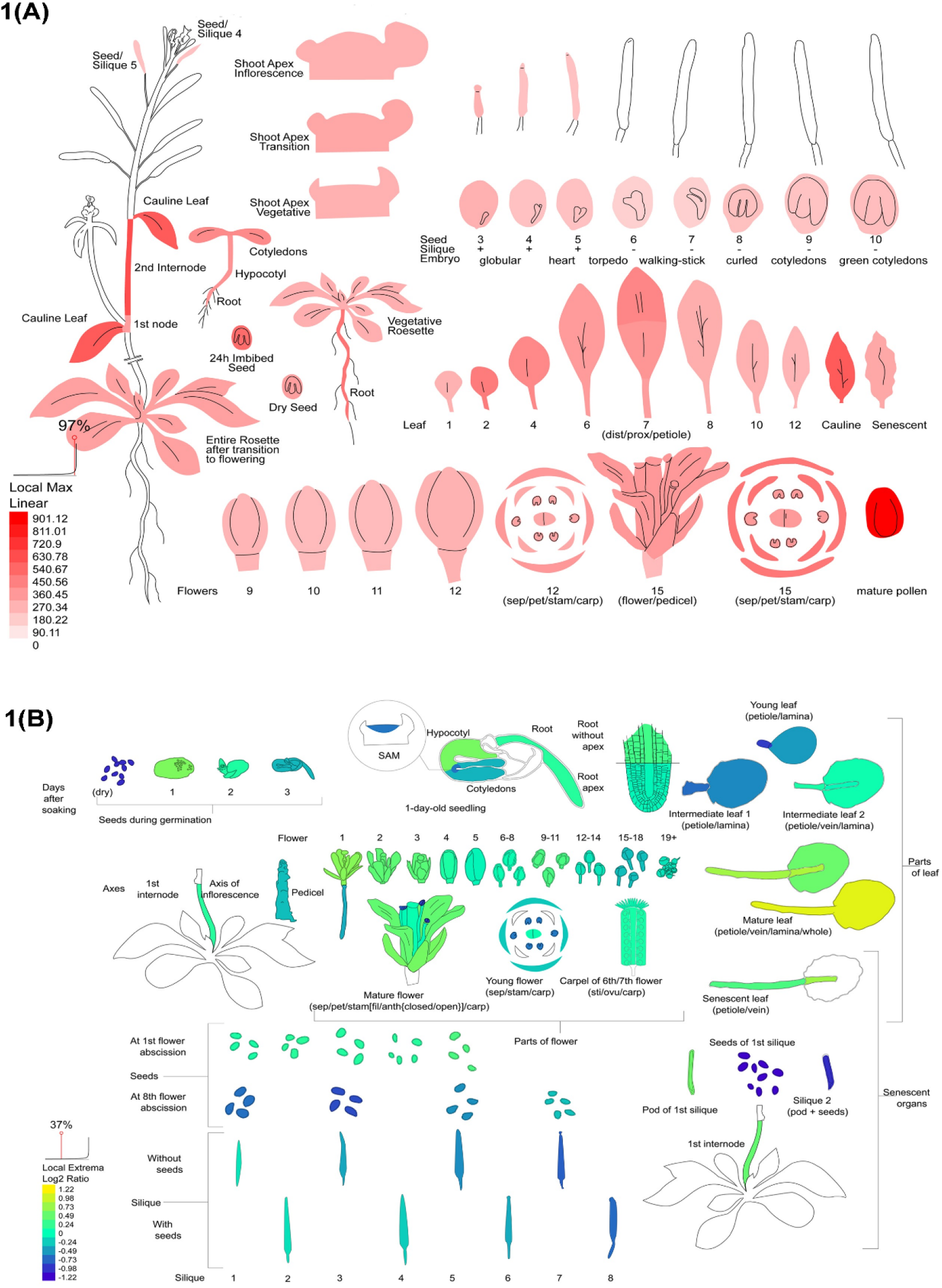
RNA-Seq data and developmental transcriptome expression. A: ACD11 gene expression in developmental transcriptomics. B: ACD11 gene expression in RNA-Seq transcriptomics.

#### 3.2.3 An insight of expression data based on different parameter comparison

We anticipated that our target gene ACD11 expresses significantly at various plant growth stages based on our microarray data. We also used prediction findings in this study to see how far the data is related to one another. According to parameter-based RNA-Seq data, the virulent bacterium infected 6-week-old short day plant leaf had the highest number of read counts in the locus with the highest total number of reads, as well as a higher rpb (point basal correlation) and RPKM value. However, the highest percentage of rpb and RPKM values are detected in 5-day old dark growing seedlings and etiolated 5-day old seedlings. The amount of readings in total are counted but read mapped per locus is not. Moreover, seedling and floral bud stages had the maximum rpb value and the lowest percentage of reads mapped and RPKM value. In addition, the highest number of reads mapped to a locus were observed in the leaves of long-day and short-day grown plants, the root tip of dark-raised seedlings, variously treated seedlings (e.g., NaCl, cytokinin, etc.), and plants infected with virulent pathogens (**Fig 2A**). Developmental transcriptome data, like RNA-Seq data, is used to construct dendrogram clustering to estimate how closely cells are expressed. The leaf-fruit cluster and the carpel-pollen cluster had the most expression similarity, according to the findings. Then the carpel-pollen cluster had the most in common with the flower pedicel, and this cluster had the most relationship with the leaf-fruit cluster (**Fig 2B**). In these procedures, all of the data forms a cluster with each other and displays their expression affinity. With the rosette leaf, the carpel-pollen cluster had the least expression.

**Fig 2.**
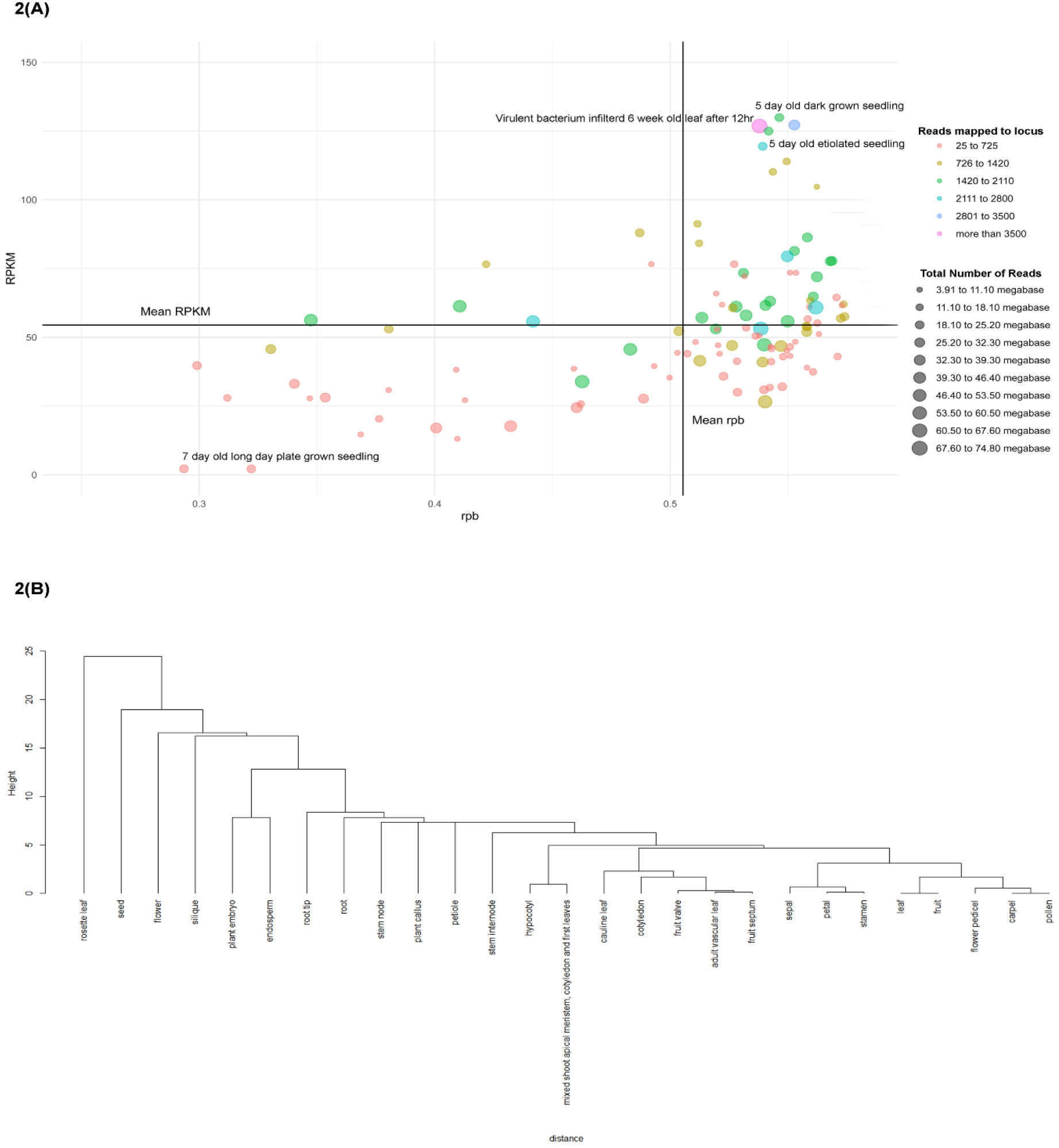
Insight of expression data based on different parameter; A: Insight on ACD11 gene data on rpb vs RPKM based on reads mapped to locus and total number of reads; B: Cluster of plant different portion based on gene expression similarity.

### 3.3 Tissue specific expression of ACD11 gene

#### 3.3.1 Gene expression in embryo developmental stage

The ACD11 gene appears to be divergent in tissue-specific embryo development. The ACD11 gene appears at every stage of embryo development, according to the microarray study. This gene expresses itself more strongly in the apical region of the globular stage than in the basal. During the embryo developing stage, the globular structure of the embryo develops into a heart shape composed of cotyledons and root. Roots express themselves significantly more effectively than cotyledons at this stage. Torpedo stage is the third stage of embryo development. It is divided into five sections: root meristem, basal, apical, and cotyledons. During the torpedo stage, the ACD11 gene exhibits itself in a unique way, with expression steadily increasing from root to cotyledons. The ACD11 gene was robustly expressed in the cotyledons during the torpedo stage, with an expression level of 2101.77. Moderate expression was observed in the apical, basal, and meristem portions, with the lowest expression predicted in the root part at 59.53 (**S6 Table and S2 Fig**).

#### 3.3.2 Gene expression in the stem epidermis and vascular bundle region

From the *Arabidopsis* microarray data analysis, we predicted that ACD11 gene expresses itself in stem and vascular bundle region. Through analysis output, it is clear that the ACD11 gene is highly expressed in the bottom portion of stem, then in the top potion and epidermal peel is expressed more strongly than whole stem. In the top portion of stem, epidermal peel was expressed negatively compared to the whole stem. On the other hand, the gene expresses itself in the bottom epidermal peel more vigorously than the whole bottom stem (**S7 Table** and **S3 Fig**). ACD11 gene expression was assessed in the cork and xylem areas in addition to the stem epidermis. We compared several genotypes of *Arabidopsis* plants in our xylem and cork expression study. Compared to Col-0 and MYB61 knockout genotypes, the ACD11 gene is substantially expressed in the cork area in the MYB50 knockout genotype, according to the study results. However, this gene was expressed more significantly in the xylem area throughout the Col-0 genotype than in the MYB61 knockout genotype, whereas MYB50 knockouts showed no expression. Different forms of expression were observed between genotypes in Hypocotyl. The ACD11 gene is highly expressed in the hypocotyl area of the plant stem in the Col-0 genotype, whereas the aba1 genotype had the lowest projected expression. The expression sequence of the ACD11 gene within different *Arabidopsis* genotypes from highest to lowest expression was observed in Col-0, axr1, max4, abi1, Ler, and aba1 genotype respectively (**S8 Table** and **S4 Fig**).

#### 3.3.3 Gene Expression in micro gametogenesis, stigma and ovaries

As RNA-Seq and developmental transcriptome data predicted that our target gene ACD11 was highly expressed in the mature pollen, so our data analysis was focused on micro gametogenesis, stigma and ovaries. From stigma and ovary analysis output, it was predicted that ACD11 gene is vigorously expressed in ovary tissues with an expression level of 634.27 and poorly expressed in stigma tissues with an expression value of 285.77 (**S9 Table** and **S5 Fig**). Apart from stigma and ovary expression analysis, we also observed expression of gene at the pollen developing stage (micro gametogenesis). The RNA-Seq and developmental transcriptome data fit seamlessly with our findings. According to the findings, the ACD11 gene is more consistently expressed in mature pollen grains than in Bicellular Pollen. The expression data demonstrated that the ACD11 gene slightly shows up in uninucleate microphore and then drops its expression in bicellular pollen. After that, it gradually intensified its expression in tricellular pollen and maximize its expression in mature pollen grain (**S10 Table** and **S6 Fig**).

### 3.4 Expression analysis of ACD11 gene in biotic and abiotic stresses

#### 3.4.1 Abiotic stress and ACD11 gene expression

When plants are subjected to biotic stressors, the ACD11 gene expresses itself. We investigated ACD11 gene expression under diverse abiotic circumstances such as heat, cold, osmotic, salt, drought, wounding, and other environmental variables. This discovery implies that, the ACD11 gene expresses itself uniquely depending on the stressor. The results from the control samples analysis suggested that this gene had not been overexposed. Different biotic stress conditions, on the other hand, predicted that the ACD11 gene was expressed both positively and negatively (**Fig 3**). This gene expressed itself highly within half an hour of being exposed to cold biotic stress, but its expression gradually declined over time. However, in the presence of osmotic stress, the ACD11 gene rapidly expressed itself within nearly an hour, then progressively decreases its expression for the next 6 hours, before gradually increasing its expression over the next 24 hours. The ACD11 gene expresses positively around half an hour of being exposed to salt, then progressively reduces its expression until it reaches 3 hours, then steadily raises its expression until it reached to 12 hours. This gene is adversely expressed for the first 3 hours of drought biotic stress, then increased its expression for the next 24 hours. When a plant is injured, the ACD11 gene expressed strongly for approximately nearly an hour and starts to increase its expression throughout the next 24 hours. When a plant is introduced to a heated environment, it expresses itself slowly for the first half hour, then gradually decreases for the next couple of hours, and then shows a high expression level after 4 hours and slightly declines over the next 24 hours (**S11 Table**).

**Fig 3.**
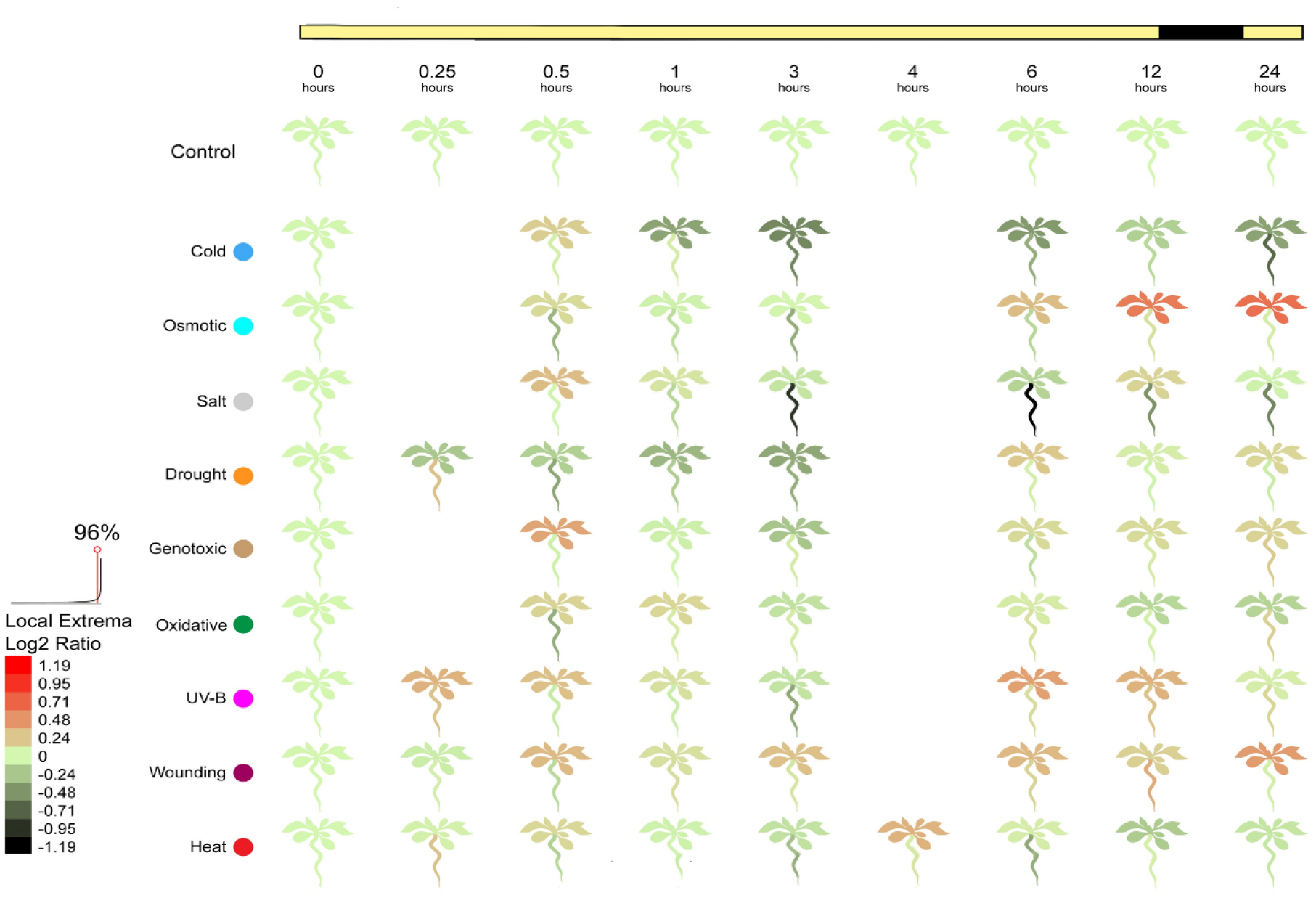
ACD11 gene expression in different abiotic stresses.

#### 3.4.2 Pathological and entomological aspect

In aspect of plant-pathogen interaction, the ACD11 gene revealed dramatically high expression when plants were subjected to any biotic stresses such as Phytophthora infestans. The experimental data predicted that when plants get afflicted by Phytophthora infestans, the expression of ACD11 elevated immensely. When half of the leaf within a plant gets affected by an avirulent pathogen Phytophthora infestans (ES4326/avrRpt2), the expression of the ACD11 gene increased slightly after 4 hours of infection,. In the next few hours, the expression dropped gradually. Subsequently, after 16 hours, the expression increased gradually up to 24 hours, then dropped slightly after 48 hours. In contrast, when the full leaf of a plant is treated with a virulent pathogen (ES4326), the ACD11 gene expression gradually increased for up to 48 hours after infection (**Fig 4 and S12 Table**). Quite apart from pathological expression, entomological quantitative analysis demonstrated that the ACD11 gene was abundantly induced when insects (Myzus persicaere) attacked *Arabidopsis* plant. The infected plant had a substantially higher expression level than the control plant, with a value of 465.98 (**S13 Table**).

**Fig 4.**
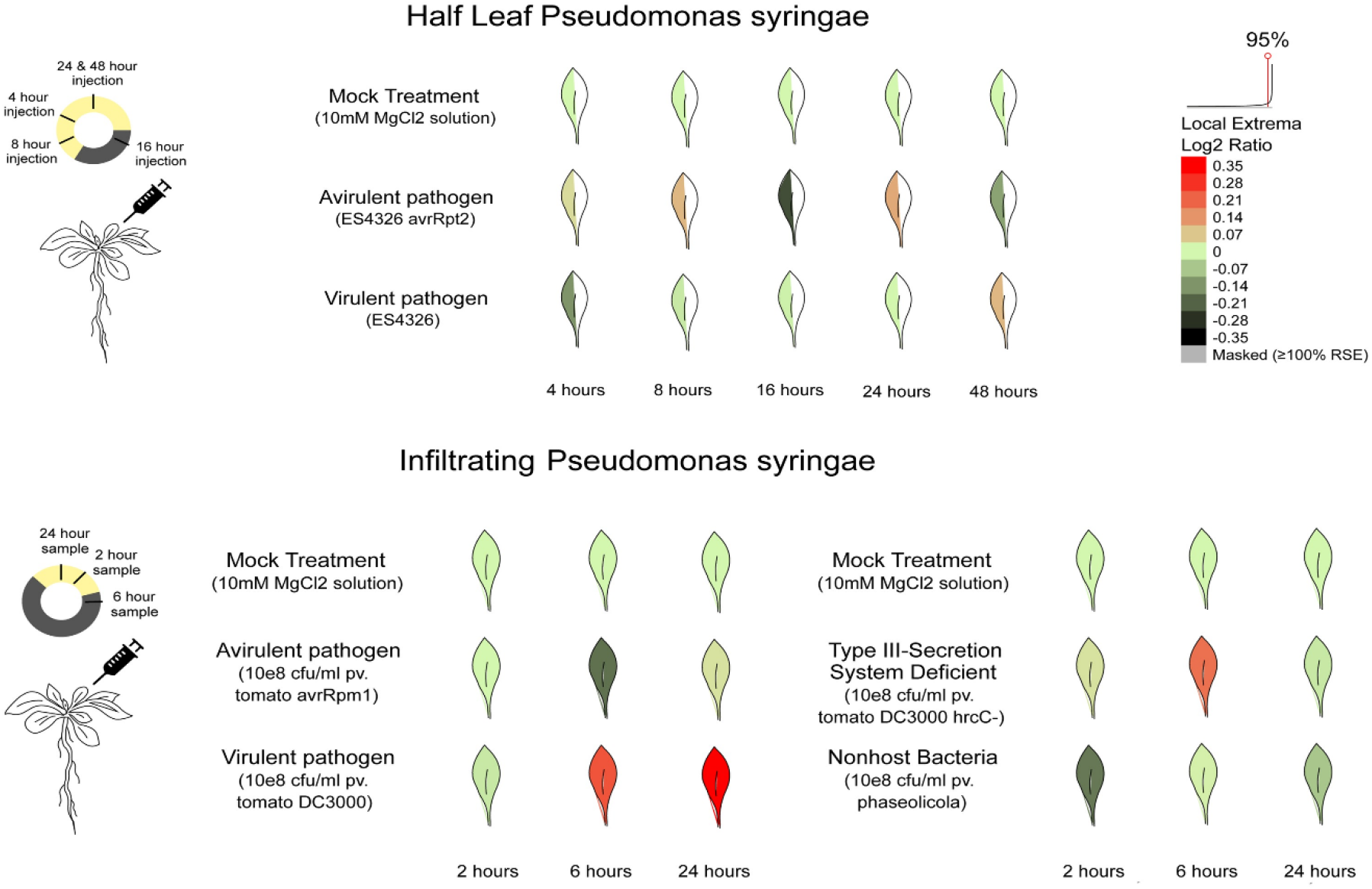
ACD11 gene expression in different biotic stresses.

### 3.5 Single Nucleotide Polymorphism (SNP) annotation in ACD11 genes

The STK11 gene polymorphism data was gathered from the 1001 genome project database, which had a total of 78 SNPs for the STK11 protein [67–68]. There were 25 SNPs in the intron area, 8 nsSNPs (missense), 4 coding synonymous, 25 in the 5′ UTR region, and 16 in the 3′ UTR region, for a total of 78 SNPs (**Fig 5**). The majority of SNPs were identified in the intron region (32.05 percent) and 5′UTR (32.05 percent), correspondingly, followed by 3′UTR SNPs (20.51 percent), missense (10.25 percent), and coding synonymous (5.13 percent). The proposed research is interested in nsSNPs because they change the encoded amino acid. For the purposes of this study, only ACD11 nsSNPs were examined (**S14Table**).

**Fig 5.**
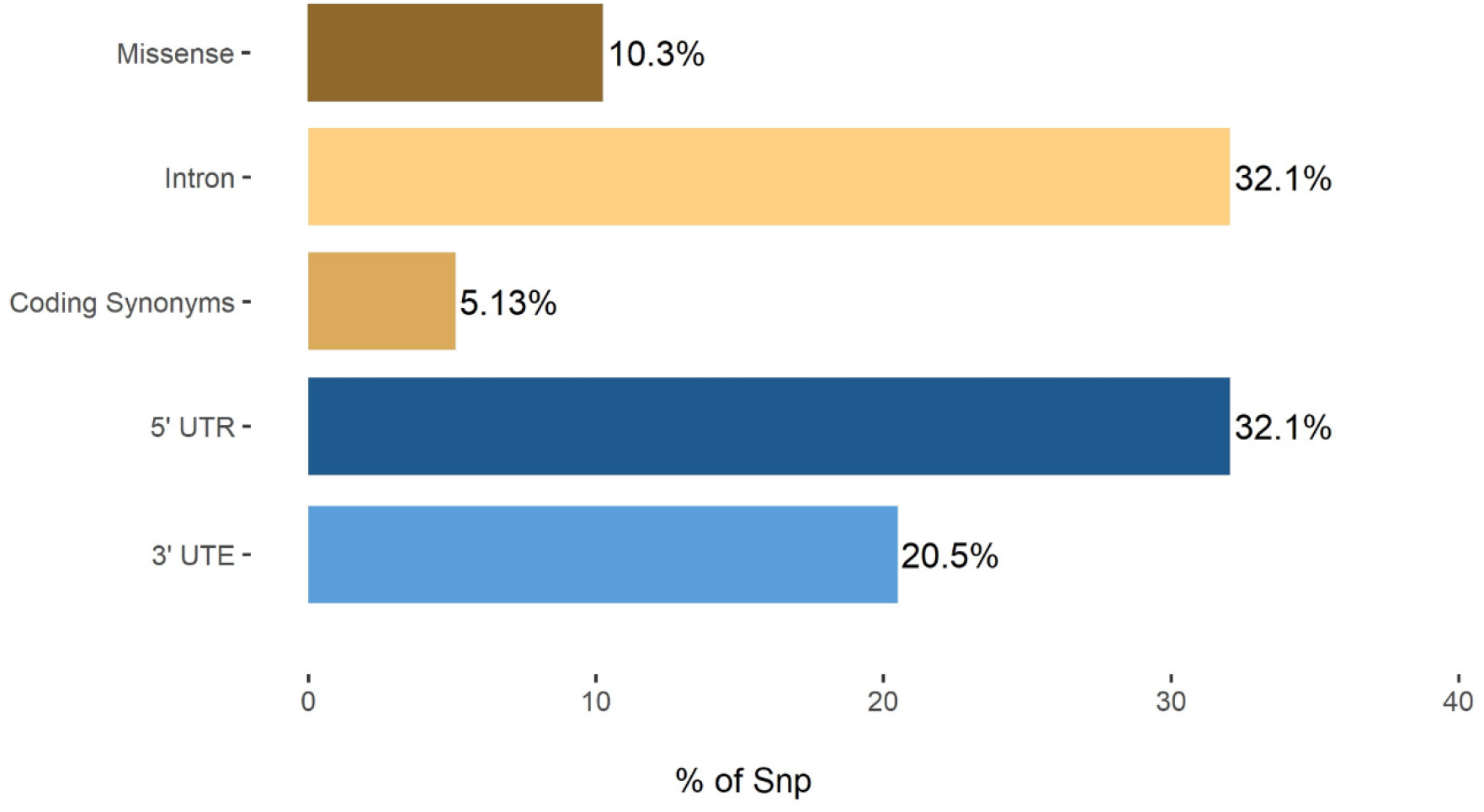
Distribution of ACD11 missense, coding synonymous, intron, 3′UTR, and 5′UTR SNPs.

### 3.6 Identification of effective SNPs in coding sequence

The aim of the numerous studies was to discover significant nsSNPs in ACD11 using computational prediction techniques. The SIFT method screened eight nsSNPs as harmful out of four missense SNPs that might have a measurable effect on the protein. Using the PolyPhen2, Panther Server, and PROVEAN algorithms, the effects of SIFT were investigated further by looking at the nsSNPs that have an impact on the structure and expression of proteins (**Fig 6**). In PolyPhen2, 3 nsSNPs were predicted to be deleterious. Panther’s evolutionary study of coding SNPs predicted 1 nsSNPs that could cause changes in protein stability due to mutation. PROVEAN anticipated that three nsSNPs were harmful and may have a practical impact on the protein. For the detection of high-risk nsSNPs in this analysis, four separate computational algorithms were used. Based on their compared prediction scores, two nsSNPs (A15T and A39D) were found to be extremely deleterious by integrating the effects of all the algorithms. A15T and A39D mutants were chosen for further investigation (**S15 Table**).

**Fig 6.**
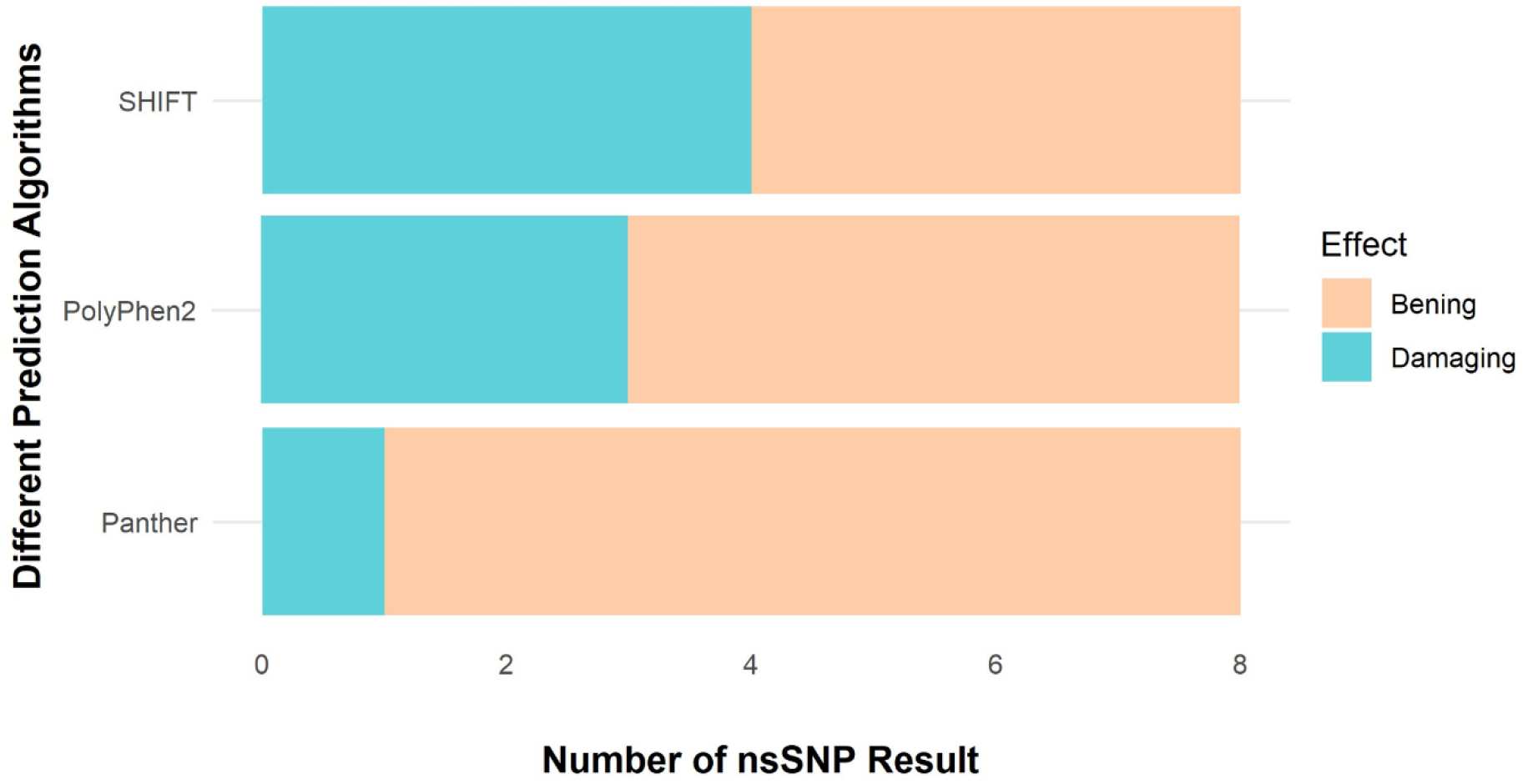
Different database data prediction.

### 3.7 Identification of potential domains in ACD11

The glycolipid transport superfamily protein ACD11 belongs to the GLTP domain-containing protein subfamily. According to previous research, this gene’s domain location varies. According to the Gene3D (1.10.3520.10) and Superfamily (SSF110004) servers, the Glycolipid transfer protein superfamily domain lies between 1-206 and 26-205 amino acids. In addition, the PANTHER (PTHR10219) and Pfam (PF08718) servers proposed that the Glycolipid transfer protein domain is placed between 5-205 and 32-169 amino acids. The chosen nsSNPs (A15T and A39D) were found in the glycolipid transfer protein domain. The glycolipid transfer protein domain contains the two nsSNPs that we looked at in this study (A15T and A39D) (**S16 Table**).

### 3.8 Structural analysis of native and mutant models

For native models, the Ramachandran plot revealed that out of 206 amino acid residue, 173 residues were in the preferred region (95.6%) and 8 residues in the allowed region (4.4%). On the other hand, the A15T mutant versions, the preferred region had 172 residues (92.0%), the approved region had 14 residues (7.5%), and the outer region had just 1 residue (0.5%). The structure assessment of A39D mutant model predicted that in the recommended zone, 166 residues (88.8%) were discovered, whereas in the allowed region, 18 amino acid residues (9.6%) were discovered. Also, there was 1 residue (0.5%) in the outer region, and just 2 residues (1.1%) in the disallowed region. Next, we considered the ERRAT and varify3D programs to determine protein structural stability and residue quality. These programs suggested that all of our native and mutant structures had extremely excellent residue coordination and backbone structures with values greater than 95% and 99.95% respectively (**S7-S9 Fig** and **S17 Table**)

### 3.9 Structural comparison of native and mutant protein

The ACD11 gene in *Arabidopsis* plays a significant function in the plant’s defense mechanism [69]. A mutation causes a substantial alteration in the protein’s structure [70]. According to our findings, the mutant form of this protein loses more interactions than the natural protein and A15T and A39D mutations trigger a significant change in the native protein structure. The alanine in position 15 has a polar interaction with the protein residues Arg11, Ser14, and Lys19 in the native structure (**Table 1**). However, when alanine is replaced with thymine in the 15th position, the protein loses the Ser14 polar interaction and gains Lys12 and Ala16 interactions (**Fig 7A**). As alanine is replaced with aspartic acid in the 39th position, the protein structure lost its Leu42 polar interaction and achieved new polar interaction with Lys19 residue (**Fig 7B**). This single point mutation has a significant influence on the overall structure of the protein. To demonstrate this point, we examined our whole protein structure and discovered that the overall number of contacts, van-der-wall interactions, polar interactions, hydrogen bonds, and ionic interactions had altered significantly (**Table 2**). Apart from this, when we super imposed our structures, we found that mutate structure build a loop where native structure had helix (**Fig 7B**).

**Fig 7.**
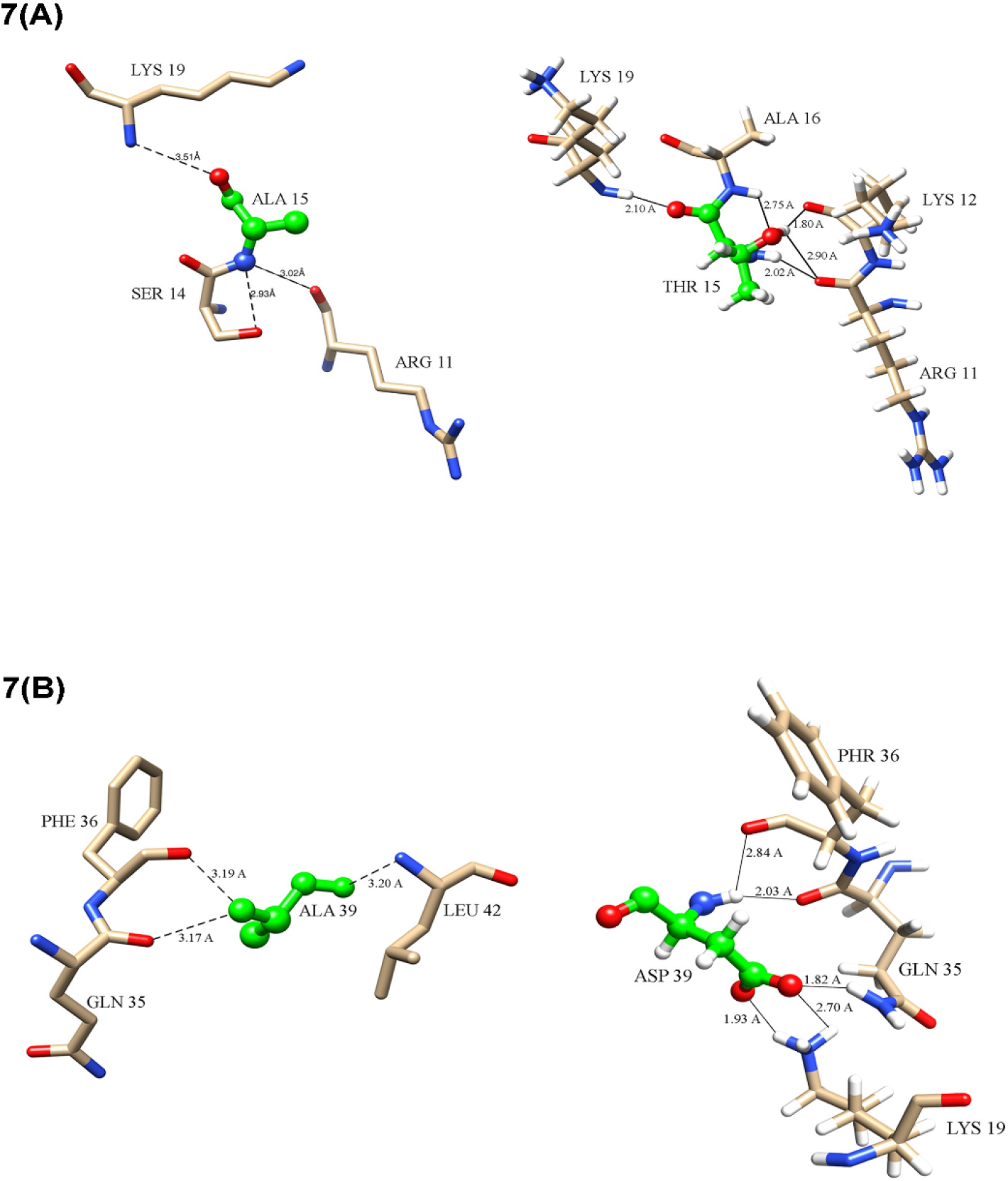
Protein ligand interaction (A) A15T mutation gained some new interaction and loses some native interaction; (B) A39D interaction also gained some new interaction.

**Table 1.** Intramolecular interactions between native and mutant protein structure (Å = 10^-10^m)

| *Interacting Residue* | *Distance ($\text{\AA}$)* | *Interacting Residue* | *Distance ($\text{\AA}$)* | *Interacting Residue* | *Distance ($\text{\AA}$)* | *Interacting Residue* | *Distance ($\text{\AA}$)* |
| --- | --- | --- | --- | --- | --- | --- | --- |
| *Native ACD11* |  | *Mutant A15T* |  | *Native ACD11* |  | *Mutant A39D* |  |
| Arg11 | 3.0 | Arg11 | 2.0 | Gln35 | 3.2 | Lys19 | 1.9 |
| Ser14 | 2.9 | Lys12 | 1.8 | Phe36 | 3.2 | Lys19 | 2.0 |
| Lys19 | 3.5 | Lys12 | 2.5 | Leu42 | 3.2 | Gln35 | 2.0 |
|  |  | Ala16 | 2.8 |  |  | Phe36 | 2.8 |
|  |  | Lys19 | 2.1 |  |  |  |  |

**Table 2.** Total number of molecular interactions of native and mutant protein.

| *Mutation* | *Total no. of Contacts* | *VdW Interactions* | *VdW clash Interactions* | *Polar Contacts* | *Hydrogen Bonds* | *Ionic Interactions* | *Aromatic Contacts* | *Hydrophobic Contacts* | *Carbonyl Interactions* |
| --- | --- | --- | --- | --- | --- | --- | --- | --- | --- |
| Native | 8084 | 188 | 277 | 411 | 191 | 24 | 30 | 488 | 7 |
| A15T | 8430 | 173 | 345 | 451 | 294 | 52 | 7 | 482 | 8 |
| A39D | 8626 | 190 | 347 | 463 | 311 | 57 | 30 | 500 | 12 |

### 3.10 Homology modelling, validation and molecular docking study

The ACD11 gene has two ligands which plays an important role in molecular activity of the gene [71]. According to the protein ligand docking review, the mutant ACD11 structure binds to the SPU and EDO ligand in a significantly different alignment than the native ACD11 structure. When compared to the A39D mutant, the A15T mutant had a greater variance. In A15T mutation, both ligand SPU and EDO binds differently than native protein structure. Besides this, the A15T mutant structure losses many of its native interactions. The native structure has binding affinity of −2.67 kcal/mol and −1.82 kcal/mol, accordingly for SPU and EDO ligands. The A15T mutant model, on the other hand, binds to SPU and EDO ligands differently, with binding affinity value of −1.15 kcal/mol and −2.65 kcal/mol, respectively. When native and mutant proteins were compared, both SPU and EDO binds to various binding pockets; however, examination of the binding pose of SPU and EDO revealed a substantial difference in both ligands’ terminal interactions between native and A15T mutant protein complexes. Certain residues in native ACD11 bind with SPU, such as Asp60 and Gly144, but these connections were lacked in mutant proteins, as Lys55 and Phe56 contacts with EDO ligand (**Fig 8**).

**Fig 8.**
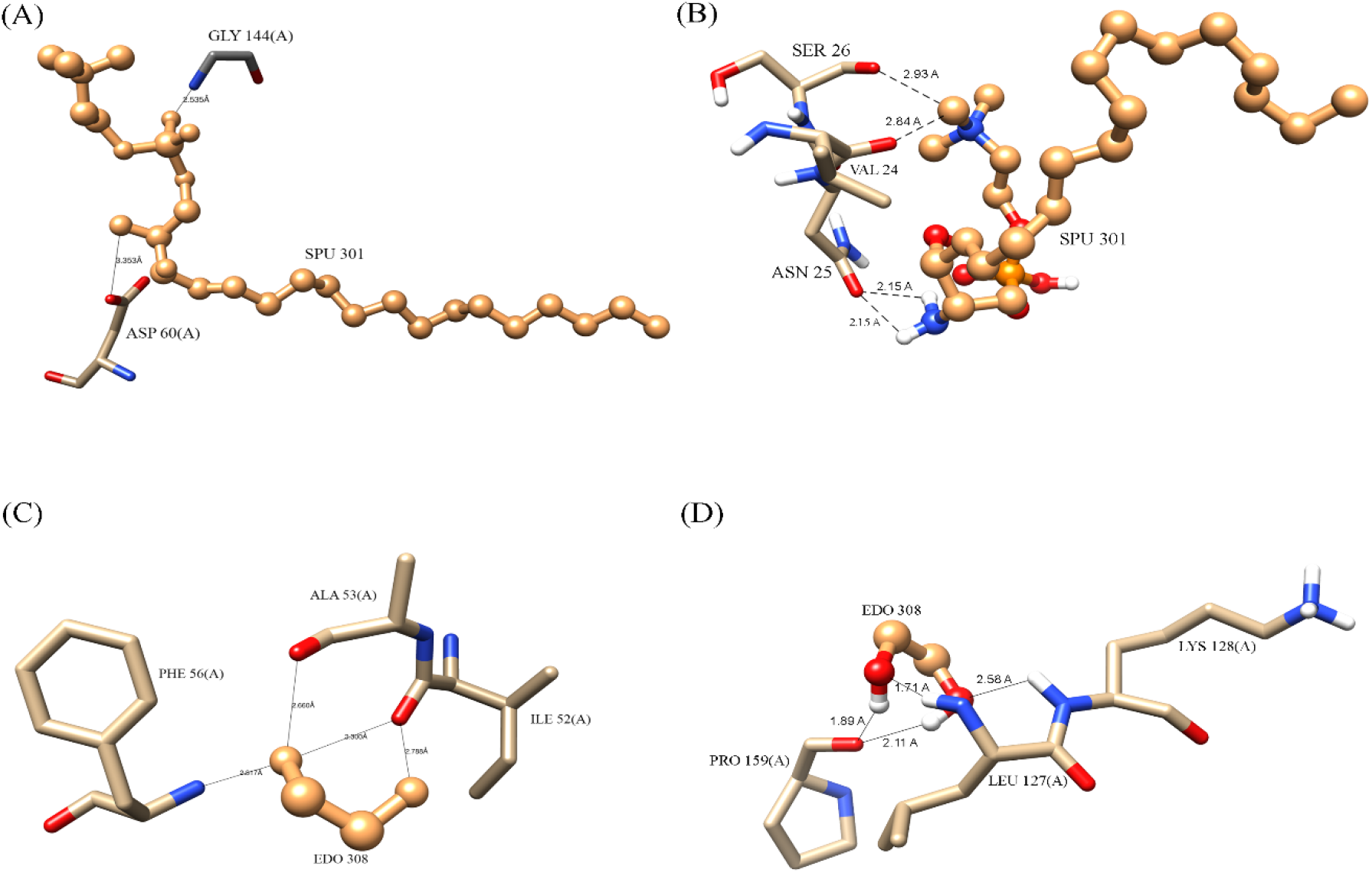
Ligand interaction change with protein structure of ACD11 because of A15T mutation.

Apart from that, the A39D mutation also causes significant differences in protein ligand binding. SPU and EDO ligands bind to the A39D mutant model with values of −1.48 kcal/mol and −2.36 kcal/mol, respectively. SPU and EDO bind to distinct binding pockets in native and mutant proteins, similar to A15T mutant structure; nevertheless, analyzing the binding posture of SPU and EDO reveals a substantial difference in the terminal contacts of both ligands between natural and mutant protein complexes. Several residues in normal ACD11 were engage with SPU, including as Asp60 and Gly144, but these interactions were absent in mutant proteins, resulting in novel associations with Thr77. Moreover, Lys55 and Phe56 contacts with EDO ligand, are missing in mutant proteins, and new interactions with Glu5 and Arg11 residue were formed (**Fig 9**).

**Fig 9.**
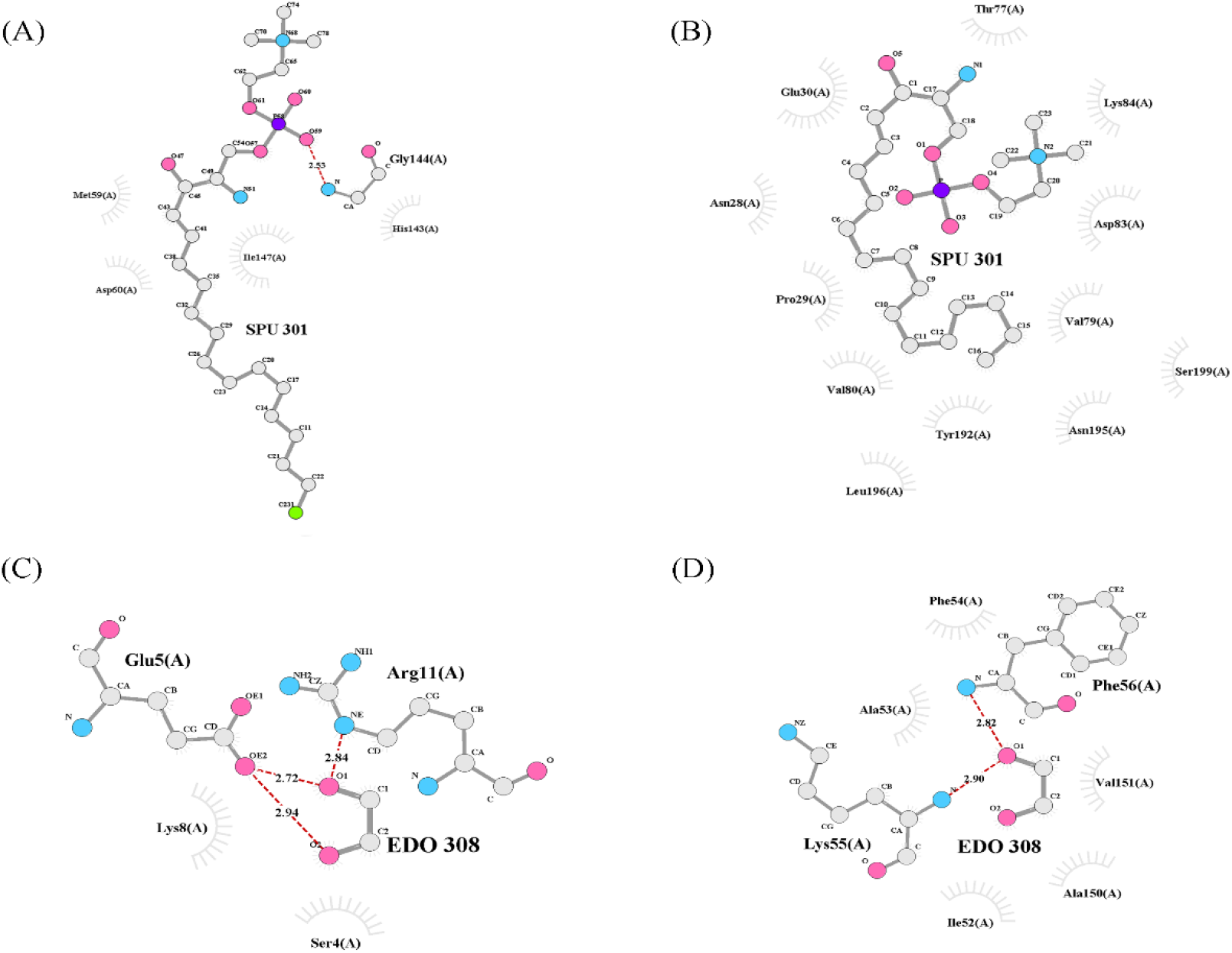
Ligand interaction change with protein structure of ACD11 because of A39D mutation.

SPU interactions with native and mutant proteins revealed less hydrogen bonds and more enticing electrostatic charge interactions between SPU and mutant protein residues whereas EDO interactions with protein residues revealed more hydrogen bonds and enticing electrostatic charge interactions in native and mutant protein structures (**Table 3**).

**Table 3:** Docking results of SPU and EDO ligands with native and mutant proteins

| No. | Compound | Protein | RMSD<sup>b</sup> | Binding<br>energy<br>(Kcal/Mol) | Inhibition<br>Constant<sup>d</sup><br>(Ki) | No of<br>H bonds | Amino acid involved<br>in interaction |
| --- | --- | --- | --- | --- | --- | --- | --- |
| 1 | SPU | Native | 43.79 | -2.67 | 11.09 mM | 2 | Asp60, Gly144 |
|  |  | A15T | 109.04 | -1.15 | 142.71 mM | 2 | Asn25, Asn25 |
|  |  | A39D | 111.73 | -1.48 | 82.37 mM | 2 | Thr77, Thr77 |
| 2 | EDO | Native | 56.13 | -1.82 | 46.01 mM | 2 | Lys55, Phe56 |
|  |  | A15T | 84.51 | -2.65 | 11.35 mM | 5 | Leu127, Lys128,<br>Pro159, Pro159,<br>Arg161 |
|  |  | A39D | 80.48 | -2.36 | 18.65 mM | 6 | Glu5, Glu5, Glu5,<br>Glu5, Arg11, Arg11 |

## 4. Discussion

The present study findings make a correlation between mutational structural changes and molecular function alteration. As plants introduce genetically mediated mechanisms such as accelerated-cell-death 11 (ACD11) for researching localized cellular suicide, and programmed cell death (PCD) for preventing pathogen dissemination throughout the plant, the recessive *Arabidopsis* mutant with accelerated cell death11 (ACD11) is identified [8]. ACD11 is a ceramide-1-phosphate (C1P) and phytoceramide-1-phosphate intermembrane transport protein [6]. ACD11 is a plant gene in *Arabidopsis thaliana* plant that induces defense-related programmed cell death (PCD), growth inhibition, and premature leaf chlorosis in seedlings before flowering, resulting in a lethal phenotype [75]. The ACD11 gene is also linked to the glycolipid transport protein family (GLTP) found in mammals [74] and enhances sphingosine transport [8].. In our ATH1 microarray data analysis, the ACD11 gene is favorably expressed in mature tissues of plants components such as cauline leaf and mature pollen, and negatively expressed in the early stages of plant growth. Moreover, ACD11 gene plays vital role in plant immunity because it prevents pathogen buildup in the plant body through constitutive defense responses [76]. As assessed by flow cytometry, ACD11 cell death is similar to mammalian apoptosis, and ACD11 produces protective genetic traits constitutively, which are linked to the hypersensitive reaction induced by virulent and avirulent pathogens [8]. Our RNA-Seq study also illustrated that the ACD11 gene was expressed robustly when *Arabidopsis* plants were continually exposed to viruses and various biotic and abiotic stressors. So, we hypothesis that deleterious mutations might have huge impact on ACD11 gene functions as well as on the structure. Therefore, to validate our assumption, we performed some *in silico* prediction analysis. We used The Project HOPE web server to calculate the evolutionary stability characteristics of all ACD11 amino acid residues in order to analyze the two nsSNPs that have a negative influence (A15T and A39D) on the ACD11 protein [77]. Alanine, at position 39, is projected to be an embedded composition and amino acid residue with a significant sustainability score by this server. This mutant residue adds a negative charge to a buried residue, perhaps results in protein folding issues. Our findings also implies that the A15T and A39D mutations alter the structure as well as amino acid interactions of ACD11 gene. For further understanding we used molecular docking analysis to test our hypothesis that the A15T and A39D mutants have a deleterious impact on the ACD11 protein. The binding pocket of ACD11 was greatly perturbed by both mutants, according to docking analysis with SPU and EDO ligands. In the native ACD11-SPU complex, SPU binds to Asp60, Gly144 but in A15T mutant-SPU complex it binds to Asn25 and same event happed with A39D mutant-SPU complex as it binds with Thr77. As a consequence, the SPU ligand binds loosely to the mutants then the native structure. In the native ACD11-EDO complex, EDO binds to Lys55 and Phe56, but in the A15T mutant-EDO complex, it binds to Leu127, Lys128, Pro159, and Arg161, and in the A39D mutant-EDO complex, it binds to Glu5 and Arg11. As a result, the EDO ligand binds with mutants of ACD11 more tightly than it does to the native protein structure. The favorable contacts needed for ACD11’s functional activity are disrupted by these mutants. It has been proven in previous studies that when a cell loses its binding affinity or interaction with SPU and increases its interactions with EDO, cell death multiplies exponentially [78–79]. In addition, SNPs in *Oryza sativa* induce seed shattering [80]. As a whole, our research indicated that our computational findings were significantly correlated with prior research results. Our study extends our knowledge of how a polymorphism impacts plant phenotypes at the molecular level. As a consideration, large-scale field experiments on a significant population are needed to classify the SNP evidence, as well as experimental mutational studies to validate the results.

## Acknowledgements

The authors would like to acknowledge Sylhet Agricultural University, Sylhet-3100 for giving the opportunity and environment to carry out the research work.

## Author Contributions

Conceptualization: Mahmudul Hasan Rifat

Data curation: Mahmudul Hasan Rifat, Jamil ahmed

Formal analysis: Mahmudul Hasan Rifat, Jamil Ahmed

Investigation: Airin Gulsan, Milad Ahmed

Methodology: Foeaz Ahmed, Milad Ahmed, Mahmudul Hasan Rifat, Jamil Ahmed

Resources: Mahmudul Hasan Rifat

Software: Mahmudul Hasan Rifat, Mahmudul Hasan, Foeaz Ahmed, Milad Ahmed, Jamil Ahmed

Supervision: Mahmudul Hasan

Validation: Mahmudul Hasan

Visualization: Mahmudul Hasan Rifat

Writing – Mahmudul Hasan Rifat, Jamil Ahmed, Airin Gulsan, Foeaz Ahmed, Milad Ahmed, Mahmudul Hasan

Writing – Mahmudul Hasan Rifat, Jamil Ahmed

## Supporting information

S1 Fig. Cellular localization.

S2 Fig. Embryo developmental data of microarray analysis

S3 Fig. Tissue specific expression of ACD11 gene in stem epidermis

S4 Fig. Tissue Specific Expression of ACD11 gene in xylem and cork

S5 Fig. Tissue specific expression of ACD11 gene in stigma and ovaries

S6 Fig. Tissue specific expression of ACD11 gene in micro gametogenesis

S7 Fig. Ramchandra plot of native protein

S8 Fig. Ramchandra plot of A15T mutant protein

S9 Fig. Ramchandra plot of A39D mutant protein

### Supplementary Table

S1 Table. Genomic data retrieval data

S2 Table. Protein information

S3 Table: Cellular localization data

S4 Table. RNA-Seq analysis data of microarray analysis

S5 Table. Developmental transcriptomics data of microarray analysis

S6 Table. Embryo developmental data of microarray analysis

S7 Table. Stem epidermis data of microarray analysis

S8 Table. Xylem and cork data of microarray analysis

S9 Table. Stigma and ovaries data of microarray analysis

S10 Table. Micro gametogenesis data of microarray analysis

S11 Table. Abiotic stress data of microarray analysis

S12 Table. Pytopthora infestance data of microarray analysis

S13 Table. Aphid infection data of microarray analysis

S14 Table. SNP Annotation

S15 Table. Functional SNP region

S16 Table. Potential domain information of ACD11

S17 Table. Ramchandra plot, ERRAT and Varify3D data

